# The PPARβ/delta-induced mesenchymal stromal cell secretome has cytoprotective effects via ANGPTL4 in a pre-clinical model of acute lung inflammation

**DOI:** 10.1101/2025.07.28.667170

**Authors:** Courteney Tunstead, Molly Dunlop, Sinéad Ryan, Evelina Volkova, Evangeline Johnston, Sabrina Batah, Claudia C. Dos Santos, Bairbre McNicholas, Claire Masterson, John G. Laffey, Karen English

**Affiliations:** Cellular Immunology Lab, Department of Biology, Maynooth University, Maynooth, Co. Kildare, Ireland; Keenan Research Centre for Biomedical Research, St. Michael’s Hospital, Toronto, Canada; Anesthesia and Intensive Care Medicine, School of Medicine, College of Medicine Nursing and Health Sciences, University of Galway, Galway, Ireland; Anesthesia and Intensive Care Medicine, Galway University Hospitals, Saolta University Hospitals Groups, Galway, Ireland; Kathleen Lonsdale Institute for Human Health Research, Maynooth University, Maynooth, Co. Kildare, Ireland

## Abstract

**Rationale:** Human bone marrow-derived mesenchymal stromal cells (hBM-MSCs) are known to exert immunomodulatory and pro-reparative effects *in vivo*. This makes hBM-MSCs an enticing therapeutic candidate for inflammatory diseases, such as acute respiratory distress syndrome (ARDS). The ARDS microenvironment is complex and contains an abundance of free fatty acids (FFAs); which are known to differentially impact MSC functionality. PPARβ/**δ** is a ubiquitously expressed nuclear receptor that is activated in response to FFA-binding. PPARβ/**δ** has been shown to impact the therapeutic efficacy of mouse MSCs.

**Objective:** This study sought to investigate the impact of PPARβ/**δ**-modulation on human MSC functionality *in vitro* and *in vivo*.

**Methods:** hBM-MSCs were exposed to a synthetic PPARβ/**δ** agonist/antagonist in the presence or absence of ARDS patient serum and the immunomodulatory and pro-reparative capacity of the MSC secretome was investigated using *in vitro* assays and a pre-clinical model of LPS-induced acute lung inflammation (ALI).

**Results:** Our results highlighted enhanced pro-reparative capacity of PPARβ/**δ**-agonised hBM-MSCs secretome in CALU-3 lung epithelial cells, mediated by MSC derived angiopoietin-like 4 (ANGPTL4). PPARβ/**δ**-induced ANGPTL4-high MSC secretome facilitated enhanced endothelial barrier integrity in the lungs of ALI mice. Therapeutic effects of PPARβ/**δ**-agonised hBM-MSCs secretome were further enhanced by licensing MSCs with human ARDS patient serum. ARDS-licensed PPARβ/**δ**-induced ANGPTL4-high MSC secretome had reduced clinical score and weight loss. The role ANGPL4 in these protective effects was confirmed using an anti-ANGPTL4 antibody.

**Conclusion:** These findings conclude that the MSC secretome therapeutic effects can be enhanced both *in vitro* and *in vivo* through licensing strategies that upregulate the angiogenic factor ANGPTL4.

**Graphical Abstract:** 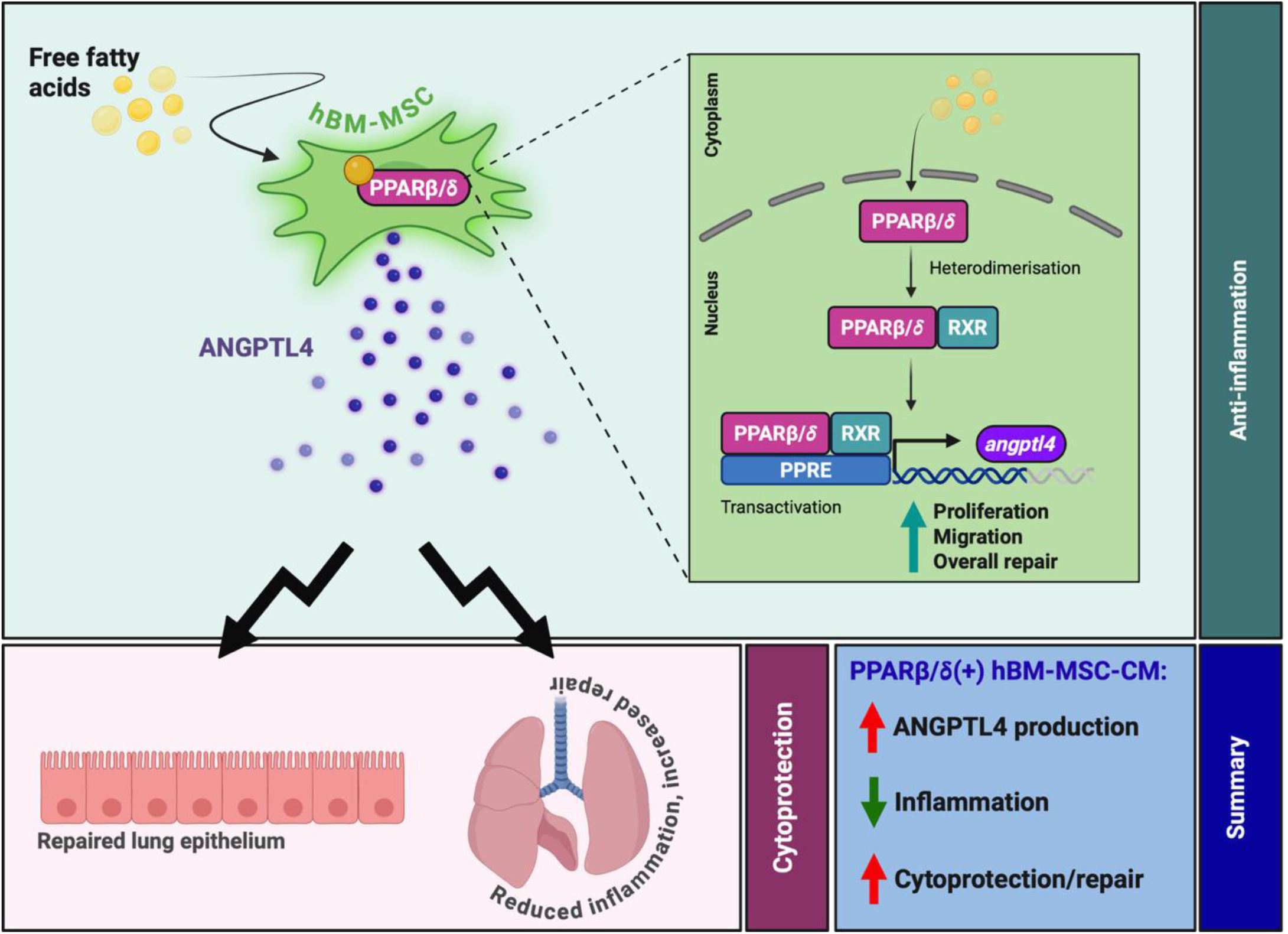

## INTRODUCTION

Cell therapy, particularly mesenchymal stromal cell (MSC) therapy, has long been thought of as a therapeutic candidate in the treatment of many inflammatory diseases. This is largely due to their immunomodulatory and cytoprotective capacity [1]. One of the conditions that MSCs have been considered for is acute respiratory distress syndrome (ARDS); an inflammatory, degenerative condition of the lung [2, 3]. MSCs have been trialled and tested for ARDS over the last 20 years, but clinical trials have shown poor outcomes; with ∼60% non-responders [4–6]. The environment into which MSCs are administered can alter their functional capacity [7–9]. There is limited understanding of how the ARDS microenvironment may impact MSCs. The complex ARDS microenvironment contains inflammatory-mediators, microorganisms, and free fatty acids (FFAs). These components have all been shown to impact MSC therapeutic capacity in various ways [10, 11]. Our previous work highlighted the differential effect of the hypo-inflammatory vs hyper-inflammatory ARDS microenvironment on MSC therapeutic efficacy with a clear correlation between high levels of inflammation, and enhanced MSC-driven repair [12]. We, and others, have also demonstrated that FFAs can influence MSC-induced immunosuppressive effects [13, 14]. In this study, we wanted to investigate the influence of FFAs present in the ARDS patient microenvironment, and the impact they have on hBM-MSCs in a pre-clinical model of acute lung inflammation (ALI).

The peroxisome proliferator-activated receptor-beta/delta (PPARβ/**δ**) is a ubiquitously-expressed nuclear receptor, present on all nucleated cells including hBM-MSCs, that is activated in response to FFA exposure [15, 16]. In recent years, PPARβ/**δ** has been shown to impact MSC therapeutic efficacy [17–27]. Interestingly, there are differential findings with some studies showing that deficiency of PPARβ/**δ** enhances MSC immunosuppressive effects [21, 22], while other studies demonstrate similar immunosuppressive effects mediated by both naive and PPARβ/**δ**-deficient mouse MSCs [24]. Studies have shown enhanced immunomodulatory capacity of T-cells by PPARβ/**δ**-inhibited mouse MSCs [21, 22]. Differentially, PPARβ/**δ** antagonism of MSCs impaired MSC therapeutic effects while priming of mouse MSCs with a PPARβ/**δ** agonist enhanced MSC cardioprotective effects *in vivo* [25]. Moreover, cardioprotective effects of MSCs were shown to be PPARβ/**δ**-dependent as the therapeutic effect was lost when PPARβ/**δ**-deficient mouse MSCs were administered [24]. These findings suggest that natural ligands of PPARβ/**δ** may have positive or detrimental effects on MSC efficacy *in vivo*. Due to the abundance of FFAs in the ARDS microenvironment, we sought to investigate the impact of PPARβ/**δ** (via synthetic agonism or antagonism) on the human BM-MSC secretome; and report the impact this had on immunomodulation and cytoprotection, in a model of ALI.

## MATERIALS AND METHODS

### hBM-MSC cell culture

Human bone marrow-derived MSCs (hBM-MSCs; RoosterBio™) were cultured in Dulbecco’s Modified Eagle Medium (DMEM) containing 10% foetal bovine serum (FBS) and 1% *Penicillin streptomycin* (cDMEM). The cells were cultured for 5 days at 37℃ + 5% CO_2_. All experiments are a representation of three independent MSC donors.

### Generation and concentration of MSC-CM

hBM-MSCs (passage 2-5) from three independent donors were seeded at a density of 1×10^5^ per well in a 6-well plate in 1ml of cDMEM and allowed to adhere. Once adhered, the cells were stimulated with 1µM of PPARβ/**δ** agonist (GW0742, Tocris™) or antagonist (GSK3787, Tocris™), as previously described [12]. 24hrs-post stimulation, the cells were washed with PBS, and the media was replaced with serum-free DMEM. This was left for a further 24hrs to allow for the generation of MSC-CM. When including ARDS serum-exposure, the agonist/antagonist were washed from the cells, and 20% ARDS serum was added for 24hrs. This was then washed away, and serum-free DMEM was added for a further 24hrs to allow for the generation of MSC-CM. All MSC-CM was harvested and stored at -20℃ for future experiments. MSC-CM was then concentrated, as required, as previously described [12].

### Gene & Protein expression

For gene expression, RNA was isolated and normalised to 100 ng/µl. cDNA was generated using the QuantBio™ cDNA Synthesis kit (as per manufacturer’s instructions). Real-time reverse transcription polymerase chain reaction (RT-PCR) was run using PerfeCta SYBR Green FastMix (QuantBio™) and pre-made Sigma™ primers (supplementary table 1). Relative expression was quantified in relation to *hprt* by calculating the 2-ΔΔCT values. For protein expression, 96-well half-area ELISA plates (COSTAR™) were coated as per manufacturers guidelines (human ANGPTL4 and VEGF (R&D™)) and run as previously described [12].

### Sequencing & metabolomic analysis

hBM-MSCs were seeded at a density of 1×10^5^ in a 6-well plate and allowed to adhere. Once attached, the cells were exposed to 1µM of PPARβ/**δ** agonist (GW0742, Tocris™) or antagonist (GSK3787, Tocris™) and incubated for 6hrs. This dose was chosen based on previous research from our group [19]. The cells were harvested in TRIzol for RNA isolation. RNA was isolated through a series of chloroform and isopropanol extractions. RNA concentration and purity was assessed using Nanodrop 2000 (Thermo Fisher Scientific) and samples were sent to Novogene™ for sequencing (GEO accession no.: GSE281162). Processed RNA sequencing datasets, provided by Novogene™, were then used to identify differentially expressed genes (DEGs) of interest. Volcano plots highlighting the most significantly enriched genes were generated using Flourish™ software. Data was also collected from publicly available RNA sequencing (GEO accession no.: GSE185263) and metabolomic datasets (Study ID: ST000042; doi: 10.21228/M8SG64). Normalised values were taken from both studies and data was plotted using GraphPad Prism™ (version 10) software.

### Scratch Assay

CALU-3 lung epithelial cells (cultured as previously described) [12] were seeded at a density of 5×10^5^ in a 6-well plate. Once 80% confluent, a single vertical scratch was made down the centre of the well using a p200 tip. The cells were then washed with PBS to remove debris, and MSC-CM was added for a period of 6 - 48hrs. The cells were then fixed with 10% formalin, before allowing to dry. Once dry, the cells were stained with crystal violet, allowed to dry, and imaged. To avoid bias, a marker was used to denote an area in the upper, central and lower regions of each scratch and images were taken from these points (supplementary figure 1). Mitomycin C (Sigma™) was added to assess the impact of proliferation at a concentration of 5 µg/ml for 2hrs prior to the scratch assay.

### Scratch Assay Analysis

Three images, taken at each of the three marked points, were taken per well; for each of the three independent MSC donors (i.e. 3 images per well x 3 MSC donors = 9 images per group). Fiji/ImageJ™ software was then used to divide each picture into four distinct quadrants in an unbiased manner (Analyse -> Tools -> Grid -> set area per point to 44444 inches squared). This grid includes the addition of 4 horizontal lines imputed on the image. Using the straight-line feature in Fiji/ImageJ™ software, four measurements were then taken per picture at each of the horizontal lines. This provided 4 measurements per picture, at unbiased points, to get a clear depiction of each percentage wound closure. The average wound closure per donor was calculated, and divided by the baseline scratch value, to get the final percentage wound closure (supplementary figure 1).

### Study cohorts and ethical approval for patient samples

Ethical approval was obtained from Galway University Medical Ethics Committee and Maynooth University Ethics Committee. All participants provided full consent.

### ARDS serum extraction

Serum samples from both male and female patients, aged 40-80 years old, were taken from patients with SARS-CoV-2-induced ARDS, from the patient pools we previously characterised and described [12].

### ANGPTL4 Neutralisation

MSC-CM was exposed to 5 µg/ml of ANGPTL4 neutralising antibody (Cell Sciences™), or 5 µg/ml IgG2aκ isotype control (Rockland Immunochemicals™). This was incubated at room temperature for 1hr prior to use. This concentration was utilised as per manufacturer guidelines for our study.

### ALI mouse model

Male and female C57BL6/J mice (Charles River), aged 8+ weeks were exposed to 2mg/kg of endotoxin (Sigma, 0001:B4 Serotype), or PBS control administered intratracheally (IT) with a total of 42 animals with sample size determined based on similar studies in the published literature. For IT administration, mice were placed in an isoflurane chamber with a steady flow of both isoflurane and oxygen for 30-45 seconds, depending on the weight of the mouse. Mice were then placed on the IT stand, and a nose cone of isoflurane was used in increments throughout the administration to ensure sufficient anaesthesia. Mice were randomized to groups according to cage mate sex and ID sequence. A single animal represents the experimental unit; and within each cage, there were mice receiving different treatments to account for potential confounders with numbers per group outlined in each figure legend. There was no specific inclusion or exclusion criteria used for this study; any animals not accounted for across all groups were due to required euthanasia prior to experimental end-point and limitation on the amount of ARDS patient serum available. Animals were identified using a numerical ID for the purpose of this study. Although information regarding groupings and treatments was available to the lead researchers, the numerical IDs were used for data harvest and data analysis to prevent bias in result interpretation. MSC-CM was prepared as described in the methods section. 500µl of PPARβ/**δ**-modulated MSC-CM, i.e. the equivalent of what would be produced from 5×10^4^ (low dose) of hBM-MSCs, was then concentrated to a volume of 30µl and administered intranasally (IN) 4hrs post-endotoxin challenge under light isoflurane anaesthesia for 15-30 seconds. An Evan’s Blue permeability assay was carried out by administration of an intravenous (IV) tail vein injection of sterile-filtered Evan’s Blue Dye (10%) 1hr prior to harvesting. The mice were sacrificed using an intraperitoneal (IP) injection of pentobarbital. Absorbance readings were taken to assess the total Evan’s Blue (620nm and 740nm) in bronchoalveolar lavage fluid (BALF) blood and whole lungs. Measurements taken from the mice included (a) pro-inflammatory cytokines screening of the BALF, (b) Evan’s Blue permeability absorbances/images, and (c) clinical scoring (based on HPRA-approved scoring methodology) and weight loss. Statistical analysis was performed using GraphPad Prism 10 software and statistical tests used have been highlighted in the figure legends of each figure. The work has been reported in line with the ARRIVE guidelines 2.0.

### Statistical Analysis

Power calculations were performed to guide sample size, and data were analyzed using GraphPad Prism 10 software. One-way ANOVA, or where appropriate a Two-way ANOVA followed by the post hoc Tukey’s multiple comparison test, was used to assess significance. For figure 2C, a t-test was carried out to identify significance in DEGs. All data are presented as mean ± SEM and the n numbers are outlined in each figure legend.

## RESULTS

### The MSC_PPARβ/**δ**(+)_ secretome enhances pro-reparative capacity in CALU-3 lung epithelial cells via migration and proliferation *in vitro*

A scratch assay in CALU-3 cells was used to investigate the impact of PPARβ/**δ** agonism (+) or antagonism (-) on MSC conditioned medium (MSC-CM) capacity to promote wound healing *in vitro* (figure 1A). Our data highlighted a significant increase in wound closure in both MSC-CM_Naive_ and MSC-CM_PPARβ/**δ**(+)_ groups (figure 1B, C). Interestingly, MSC-CM_PPARβ/**δ**(-)_ promoted wound healing when compared to control medium but did not significantly enhance CALU-3 wound closure. While there was no statistical difference between MSC-CM_Naive_ and MSC-CM_PPARβ/**δ**(-)_ (figure 1B), the percentage wound closure in the MSC-CM_PPARβ/**δ**(-)_ group was less than in the MSC-CM_Naive_ group. There is a possibility that PPARβ/**δ** antagonism could lead to the production of inhibitory factors in the conditioned medium, however, our RNA-Seq data set did not identify any known inhibitory factors (see figure 2, table 2). It is known that wound closure can involve proliferation and or migration [28, 29], thus this experiment was repeated using Mitomycin C; a known proliferation inhibitor [30]. While the addition of Mitomycin C significantly reduced wound closure in the MSC-CM_PPARβ/**δ**(+)_ group at the 48hr time point, the percentage wound healing in the presence of Mitomycin C remained higher than the medium control group suggesting that both proliferation and migration may have a combination effect (figure 1D). Interestingly, significantly more wound repair was observed in the MSC-CM_PPARβ/**δ**(+)_ group compared to the MSC-CM_Naive_ group at 48hrs, while the MSC-CM_PPARβ/**δ**(-)_ group did not inhibit or enhance would healing (figure 1D). *TOP2A is* a gene largely indicative of cell proliferation [31] and has previously been used as an indicator for cell proliferation in wound healing. Gene expression analysis of *TOP2A* in CALU-3 cells highlighted a significant difference between the MSC-CM_PPARβ/**δ**(+)_ and MSC-CM_PPARβ/**δ**(-)_ groups, with the MSC-CM_PPARβ/**δ**(+)_ group having significantly more proliferative capacity than the MSC-CM_PPARβ/**δ**(-)_ group (figure 1E). Importantly, Mitoymcin C significantly reduced *TOP2A* expression in the MSC-CM_Naive_ and MSC-CM_PPARβ/**δ**(-)_ groups (figure 1E).

**Figure 1:**
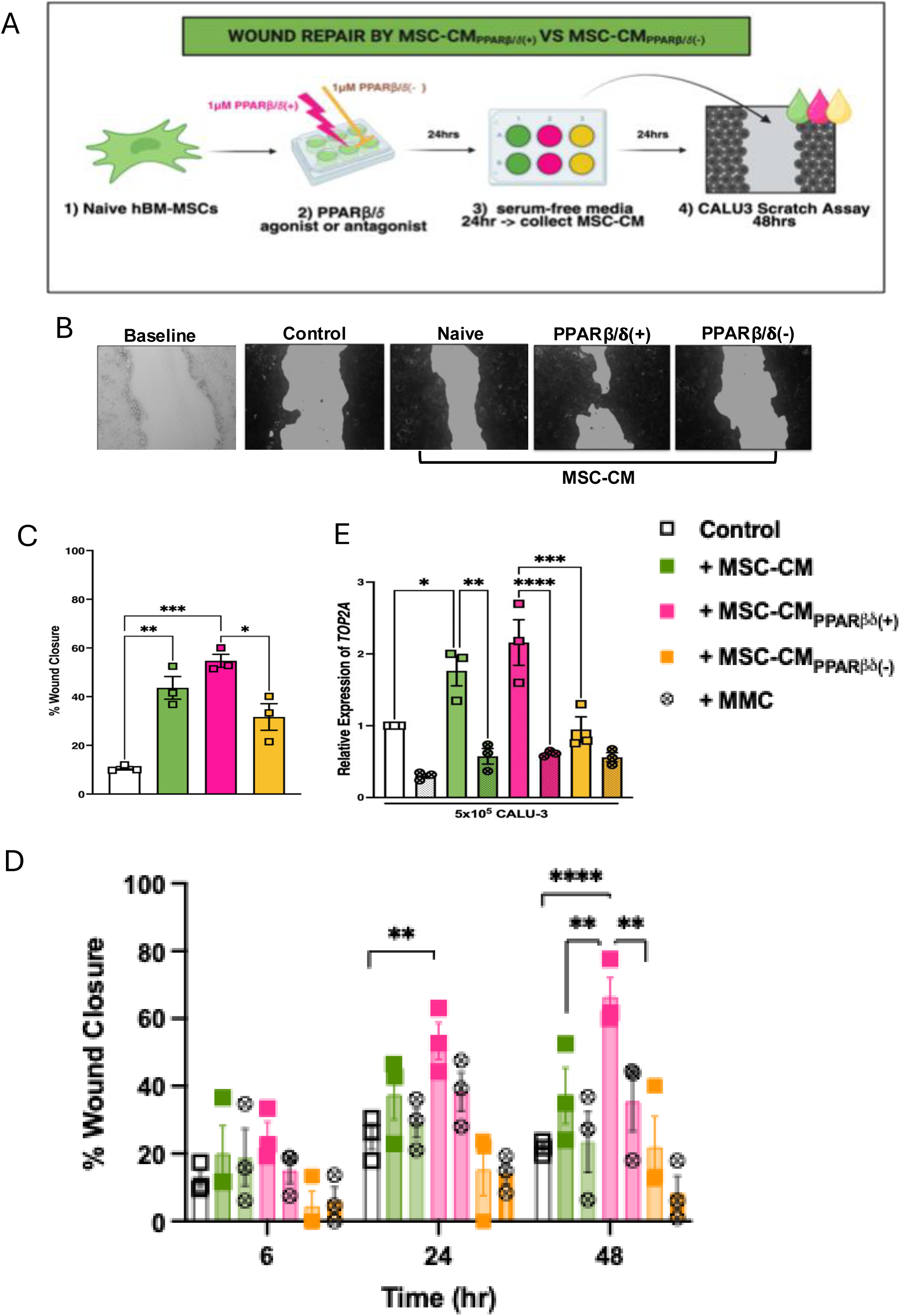
Secretome from PPARβ/**δ** agonised hBM-MSCs enhanced wound healing in CALU-3 lung epithelial cells. (A) Experimental Design. A CALU-3 scratch assay was run, in combination with MSC-CM from naive, PPARβ/**δ** agonized (+) or antagonized (-) hBM-MSCs. (B) Images of the wound healing assay at baseline or following 48hrs fixed and stained with crystal violet dye and (C) quantified using ImageJ software (one-way ANOVA followed by Tukey’s post-hoc test, n=3). White bar is CALU-3 cells in media alone. (D) Percentage (%) wound healing in scratched CALU-3 cells was recorded at different time points (6, 24 and 48hrs). Mitomycin C (MMC, denoted as circular data points) was added to CALU-3 cells 2hr prior to the scratch assay. Conditioned medium from naive, PPARβ/**δ** agonized (+) or antagonized (-) hBM-MSCs was added immediately after the scratch. Wound closure (%) was quantified using imageJ software (two-way ANOVA followed by Tukey’s post-hoc test, n=3). (E) Expression of the cell proliferation gene *TOP2A* was analysed by RT-PCR at a 48hr time-point (one-way ANOVA followed by Tukey’s post-hoc test, n=3). Replicates are a representation of 3 individual hBM-MSC donors. Data is presented as mean ± SEM; *p<0.05, **p<0.01, ***p<0.001, ****p<0.0001.

### Analysis of MSC_PPARβ/**δ**(+)_ RNA sequencing highlighted upregulation of ANGPTL4

To try to identify the potential mechanisms associated with enhanced wound healing capacity in MSC_PPARβ/**δ**(+),_ RNA sequencing (Novogene™; GEO accession no.: GSE281162) was performed on MSC_PPARβ/**δ**(+),_ MSC_PPARβ/**δ**(-)_ and MSC_Naive_ (figure 2A). The data highlighted a variety of differentially expressed genes (DEGs) between the three groups. The DEGs numbers were visualised using a Venn diagram (figure 2B), showing 170 DEGs specific to the MSC_Naive_ group, 128 specific to the MSC_PPARβ/**δ**(+)_ group, and 189 specific to the MSC_PPARβ/**δ**(-)_ group. Volcano plot visualisation highlighted several of the most significantly altered DEGs in each group (figure 2C). The MSC_PPARβ/**δ**(+)_ group had a significant enrichment of genes such as *PLIN2, ANGPTL4, FAM156A, CPT1A, CAT, NADK2, TMED6,* and *LDHAL6B.* This group also showed significant downregulation in genes such as *MAGED4* and *FAM157B* (table 1). The MSC_PPARβ/**δ**(-)_ group had a significant enrichment of genes such as *IKBKGP1, SNRPGP2, LINC01159, MIR3648-1, RARRES1, MALAT1* and *RNF217-AS1.* This group also showed significant downregulation in genes such as *RGPD5, PFN1P3, MIR3936HG, U2AF1* and *TSNAXIP1* (table 2). Fragments per kilobase of exon per million mapped fragments (FPKMs) of two known PPARβ/**δ** target genes (angiopoietin like protein 4 (ANGPTL4) and pyruvate dehydrogenase kinase 4 (PDK4)) were plotted to confirm the data was as expected (figure 2D, E), and this was confirmed by qRT-PCR (figure 2F, G).

**Figure 2:**
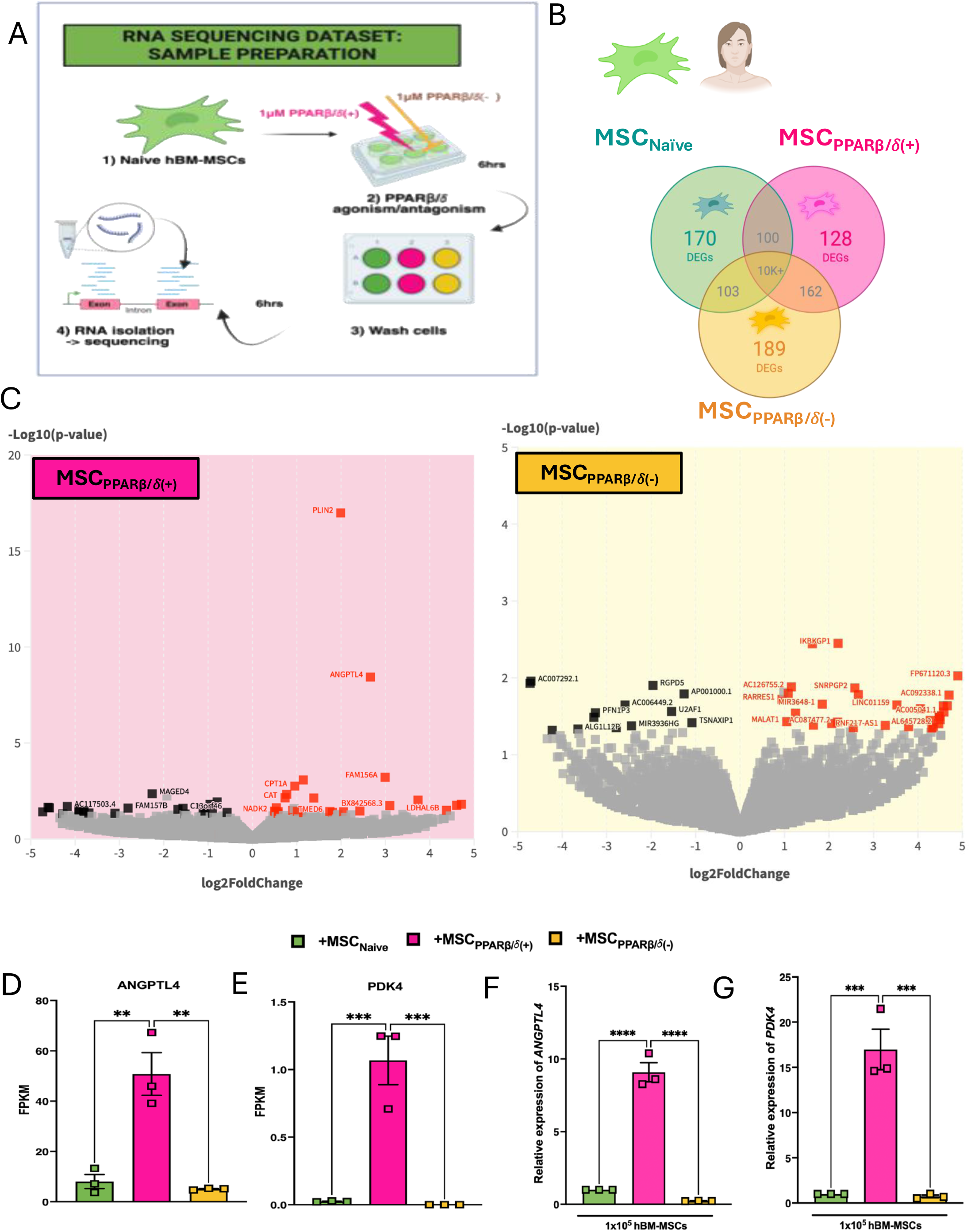
RNA-Seq identified enrichment of ANGPTL4 in PPARβ/**δ** agonised hBM-MSCs. (A) Experimental design: hBM-MSCs exposed to 1uM of PPARβ/**δ** agonist or antagonist were sent to Novogene™ for RNA sequencing. (B) Venn diagrams showing the number of differentially expressed genes (DEGs) in naive, PPARβ/**δ** agonized (+) or antagonized (-) hBM-MSCs. (C) Volcano plots generated using Flourish™ software highlight the most significant DEGs Log10(p-value). Fragments per kilobase of transcript per million mapped reads (FPKM) was plotted for two known PPARβ/**δ** target genes in the sequencing dataset: (D) *ANGPTL4* and (E) *PDK4*. Expression of *ANGPTL4* (F) and *PDK4* (G) in hBM-MSCs exposed to 1uM of PPARβ/**δ** agonist or antagonist for 6hrs was confirmed by real-time RT-PCR (one-way ANOVA followed by Tukey’s post-hoc test, n=3). Replicates are a representation of 3 individual hBM-MSC donors. Data is presented as mean ± SEM; **p<0.01, ***p<0.001, ****p<0.0001.

**Table 1:** DEG functions in MSC_PPARβ/**δ**(+)_ group.

| Gene name | Up/Down | Involved in |
| --- | --- | --- |
| *plin2* | Up | Lipid metabolism |
| *angptl4* | Up | Lipid metabolism and angiogenesis |
| *fam156A* | Up | Enables protein binding |
| *cpt1A* | Up | Lipid metabolism |
| *cat* | Up | Protection against oxidative stress |
| *nadk2* | Up | Protection against oxidative stress |
| *tmed6* | Up | Trafficking and transport |
| *ldhal6B* | Up | Lactate metabolism |
| *maged4* | Down | Cell cycle regulation |
| *fam157B* | Down | Non-coding |

**Table 2:** DEG functions in MSC_PPARβ/**δ**(-)_ group.

| Gene name | Up/Down | Involved in |
| --- | --- | --- |
| *ikbkgp1* | Up | NFκB signalling |
| *snrpgp2* | Up | RNA processing |
| *linc01159* | Up | Cell differentiation |
| *mir3648-1* | Up | Cell differentiation |
| *rarres1* | Up | Cell differentiation & retinoid signalling |
| *malat1* | Up | Cell differentiation |
| *rnf217-as1* | Up | Cell differentiation |
| *rgpd5* | Down | GTPase regulation |
| *pfn1p3* | Down | Cell motility/migration |
| *mir3936hg* | Down | Proliferation and apoptosis |
| *u2af1* | Down | ROS production/oxidative stress |
| *tsnaxip1* | Down | RNA processing and transport |

### MSC-associated genes/proteins are unaffected by PPARβ/**δ** agonism or antagonism; except ANGPTL4

Given the functional enhancement of MSC-CM_PPARβ/**δ**(+)_ promotion of wound healing observed in figure 1, the production of known cytoprotective factors released by MSCs were assessed. While MSC_Naive_ produced ANGPTL4, VEGF and MIF, MSC_PPARβ/**δ**(+)_ only produced significantly higher levels of ANGPTL4 (figure 3A-C). This aligns with the RNA-sequencing data as *ANGPTL4* was the only cytoprotective factor differentially expressed in MSC_PPARβ/**δ**(+)._ Along with this, PPARβ/**δ** agonism/antagonism did not affect expression of typical MSC-associated immunomodulatory factors such as *PTGS2, IL-6, IDO, PTGES* or *TGFβ (figure 3D-H)*.

**Figure 3:**
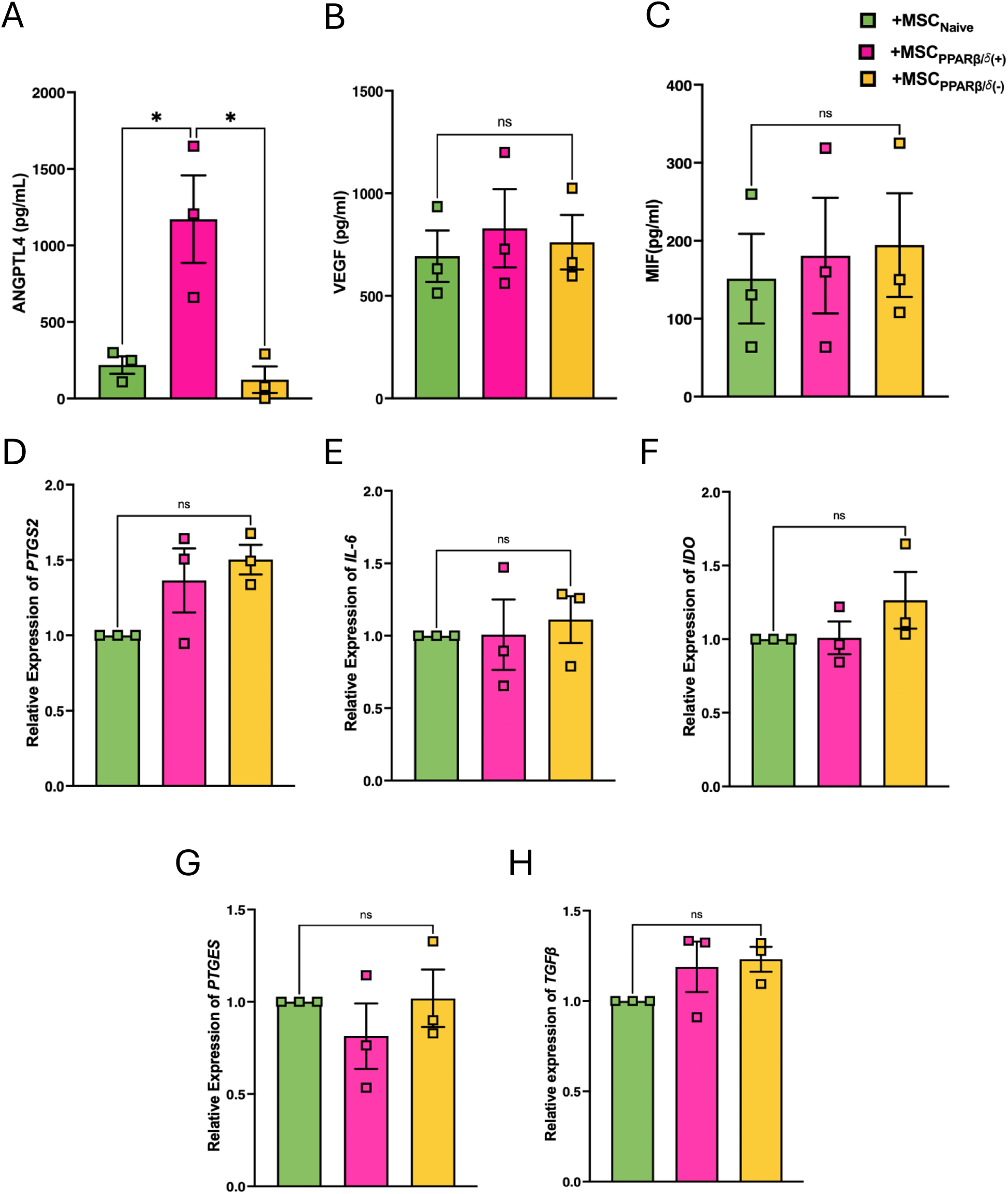
PPARβ/**δ**(+) hBM-MSCs show enhanced expression of ANGPTL4, but not other traditional MSC-associated genes or proteins. Supernatant from hBM-MSCs exposed to 1uM of PPARβ/**δ** agonist or antagonist were harvested before cells were collected in TRIzol for gene expression studies. Protein expression of (A) ANGPTL4, (B) VEGF and (C) MIF were assessed by ELISA. Gene expression of (D) *PTGS2,* (E) *IL-6,* (F) *IDO,* (G) *PTGES* and (H) *TGFβ* were assessed by real-time RT-PCR (one-way ANOVA followed by Tukey’s post-hoc test, n=3). Replicates are a representation of 3 individual hBM-MSC donors. Data is presented as mean ± SEM; *p<0.05.

### The secretome from MSC-CM_PPARβ/**δ**(+)_ enhances wound healing in CALU-3 lung epithelial cells in an ANGPTL4-dependent manner

Other studies have highlighted the role of a PPARβ/**δ** agonist [18], and ANGPTL4 in promoting repair in other tissues [29, 32]. It has also been shown that ANGPTL4 can play a significant role in proliferation and migration [28]. To determine if the increased levels of ANGPTL4 present in MSC-CM_PPARβ/**δ**(+)_ played a role in the enhanced pro-reparative capacity, we used an anti-ANGPTL4 neutralising-antibody to eliminate the functional capacity of ANGPTL4 in the MSC-CM (figure 4A). Importantly, ANGPTL4-neutralisation but not the isotype control antibody, abrogated the enhanced wound closure observed in the MSC-CM_PPARβ/**δ**(+)_ group (figure 4B-C). Addition of the ANGPTL4 neutralising antibody interfered with ANGPTL4 protein levels as quantified by ELISA (figure 4D).

**Figure 4:**
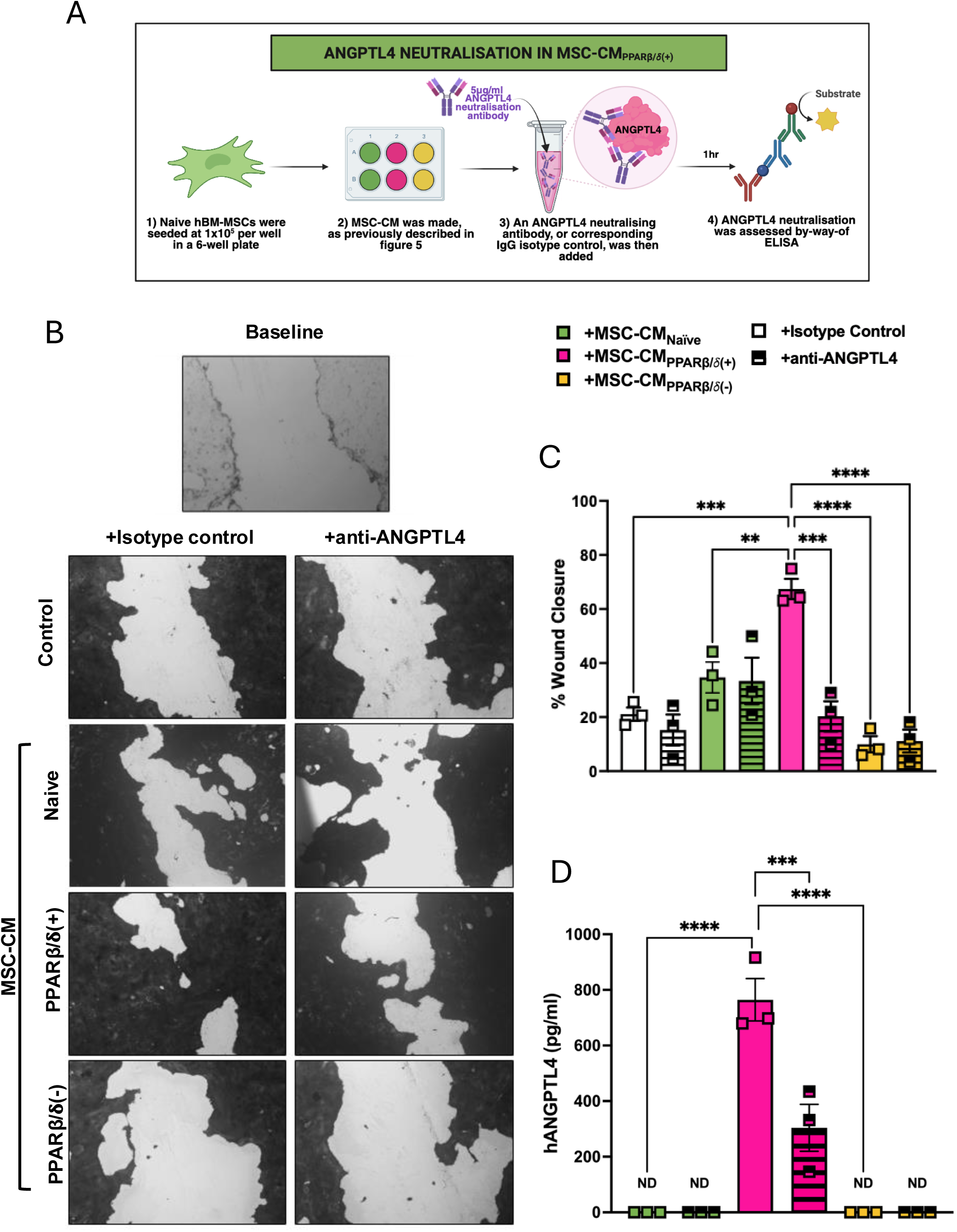
The secretome from PPARβ/**δ** agonised hBM-MSCs mediates enhanced wound repair via ANGPTL4. (A) Experimental design. Using an ANGPTL4 neutralisation approach, we assessed wound closure of CALU-3 lung epithelial cells. (B) Images taken at baseline or 48hr post scratch. Scratched CALU-3 cells were incubated with isotype control or anti-ANGPTL4 antibody before addition of conditioned medium. Medium only served as the control. (C) % wound closure quantified using ImageJ. (D) Levels of ANGPTL4 produced by naive hBM-MSCs or exposed to 1uM of PPARβ/**δ** agonist or antagonist in the presence of the isotype control of anti-ANGPTL4 antibody were measured by ELISA (one-way ANOVA followed by Tukey’s post-hoc test, n=3). Replicates are a representation of 3 individual hBM-MSC donors. ND denotes not detected. Data is presented as mean ± SEM; **p<0.01, ***p<0.001, ****p<0.0001.

### Ligands and co-activators of PPARβ/**δ** are increased, while co-repressors of PPARβ/**δ** are decreased, in ARDS patients

Thus far, our data has shown that PPARβ/**δ** agonism leads to enhanced cytoprotective effects of MSC-CM. Previously, it was deduced that MSC-CM derived from ARDS-serum licensed MSCs has superior therapeutic effects in a pre-clinical model of ALI [12]. To ensure translational relevance of our findings from this study in human conditions such as ARDS, a publicly available metabolomics dataset (Study ID: ST000042; doi: 10.21228/M8SG64) was mined. From this metabolomics dataset we identified that PPARβ/**δ** ligands are more prevalent in ARDS. Higher levels of linoleate, palmitoleic acid and arachidonate were present in the bronchoalveolar lavage fluid (BALF) from ARDS patients, in comparison to healthy controls (figure 5A). Using a publicly available RNA sequencing dataset (GEO accession no.: GSE185263), it was shown that several PPARβ/**δ**-associated co-activators that were increased in the blood of ARDS patients when compared to Non-ARDS controls (figure 5B). This included a significant increase in *NCOA1* and *NCOA3,* a trend toward increased *PPARGC1A* and *NCOA2,* and no difference in *CREBBP* (figure 5C-G). Along with this, several PPARβ/**δ**-associated co-repressors were decreased in the blood of the same ARDS patients (figure 5B). This included a significant reduction in *SIRT1, NCOR1* and *NCOR2* (figure 5H-J). Given that *ANGPTL4* was significantly upregulated in response to PPARβ/**δ** agonism in our own sequencing dataset (figure 2), levels of ANGPTL4 were further assessed upon MSC-exposure to ARDS patient serum. In comparison to the PPARβ/**δ** agonist, exposure of MSCs to ARDS patient serum significantly increased MSC secretion of ANGPTL4. Addition of both the PPARβ/**δ** agonist and ARDS patient serum further enhanced MSC production of ANGPTL4 (figure 5K). Interestingly, addition of the PPARβ/**δ** antagonist could not block the ARDS serum induction of ANGPTL4 secretion by MSCs. This is largely down to PPARβ/**δ** having multiple binding-sites [33], whereby both agonists and antagonists can bind simultaneously; with agonism out-competing antagonism (supplementary figure 2).

**Figure 5:**
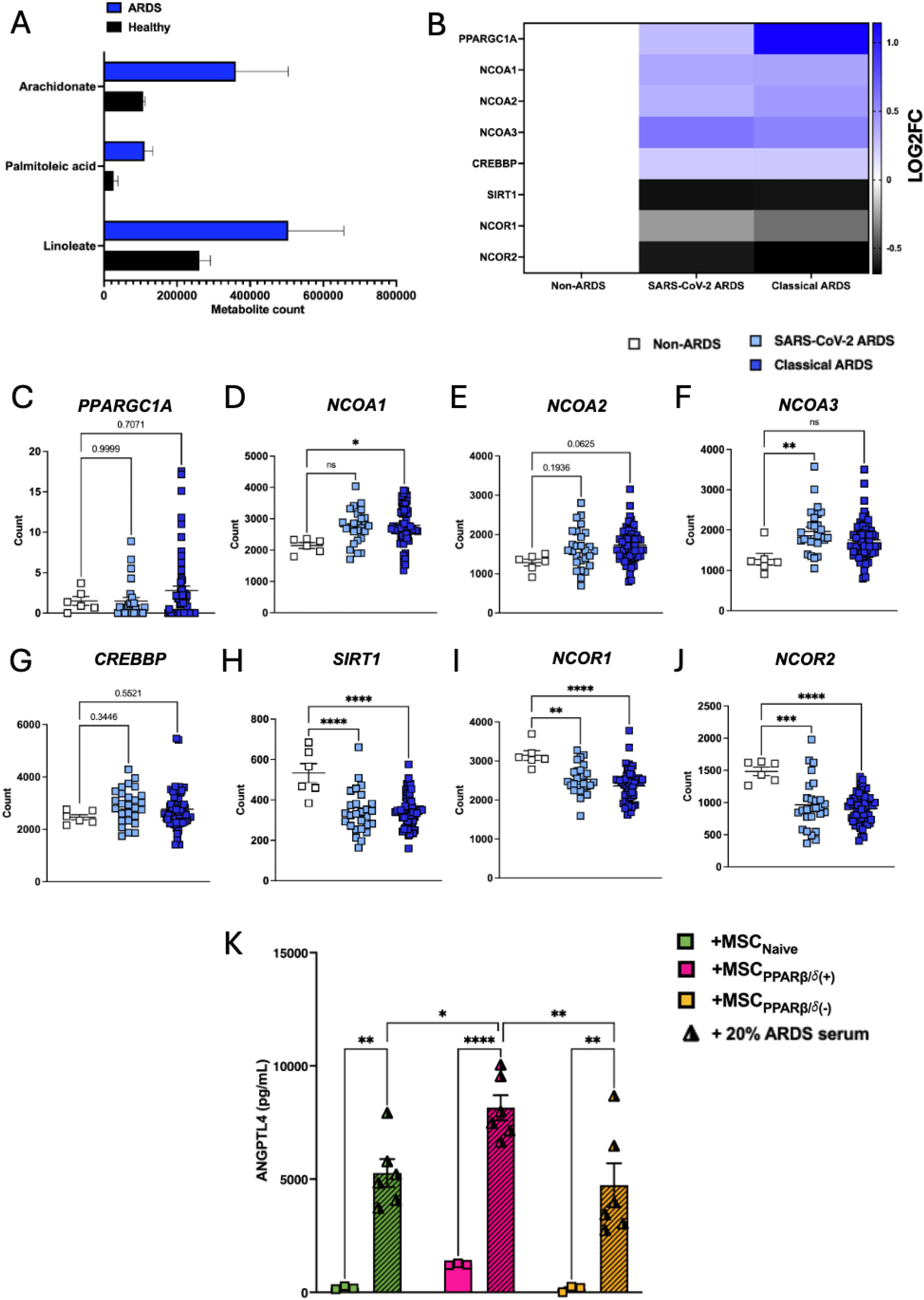
ARDS patients have higher levels of PPARβ/**δ** ligands and co-activators. (A) A publicly available dataset (Study ID: ST000042; doi: 10.21228/M8SG64) was mined for PPARβ/**δ** ligands present in the ARDS lung. Levels of Arachidonate, Palmitoleic acid and Linoleate were elevated in ARDS bronchoalveolar lavage fluid compared to healthy controls. (B) An additional RNA-Seq dataset (GEO accession no.: GSE185263), based on ARDS serum, highlighted an increase in PPARβ/**δ**-associated co-activators in both SARS-CoV-2 ARDS and Classical ARDS patient serum compared to Non-ARDS controls. RNA-Seq data counts for PPARβ/**δ**-associated co-activator genes (C) *PPARGC1A*, (D) *NCOA1*, (E) *NCOA2*, (F) *NCOA3* and (G) *CREBBP;* and co-repressor genes (H) *SIRT1*, (I) *NCOR1* and (J) *NCOR2* were plotted comparing expression between SARS-CoV-2 ARDS, Classical ARDS and Non-ARDS controls (one-way ANOVA followed by Tukey’s post-hoc test, n=6-58). Each data point (A-J) is from an individual ARDS patient, or healthy control. (K) ANGPTL4 production from 3 individual hBM-MSC donors exposed to PPARβ/**δ** agonist (+) or antagonist (-) in the presence or absence of 20% ARDS patient serum was measured by ELISA. Data is presented as mean ± SEM; *p<0.05, **p<0.01, ***p<0.001, ****p<0.0001.

### MSC-CM_PPARβ/**δ**(+)_ provides enhanced lung endothelial barrier function in ALI

Next, we investigated the protective effects of MSC-CM_PPARβ/**δ**(+)_ in a preclinical model of LPS induced-ALI. First we set out to determine the number of MSCs that could provide a therapeutic effect in a mouse model of ALI, comparing two different doses of MSCs. Administration of high dose MSCs (1×10^5^) led to significant decrease in TNFα, IL-1β and IL-6 in the BALF of ALI mice, while low dose MSCs (5 x 10^4^) could only decrease TNFα and led to increased IL-6 in the BALF (supplementary figure 3). Following this, MSC-CM that was generated from the equivalent of the low dose was used; with the idea that using a low dose would allow us to detect any enhanced therapeutic efficacy following PPARβ/**δ** agonism or antagonism. This showed that MSC-CM_PPARβ/**δ**(+)_ but not MSC-CM_PPARβ/**δ**(-)_ significantly improved the therapeutic efficacy of the non-efficacious dose of MSC-CM. Lungs of mice that received MSC-CM_PPARβ/**δ**(+)_ exhibited enhanced vascular and epithelial barrier function measured using Evans Blue dye administration (figure 6A, B). Although modest, this result was statistically powered. This effect was somewhat impacted by pre-exposing the MSC-CM to anti-ANGPTL4 neutralising antibody, but this was not significantly different from the MSC-CM_PPARβ/**δ**(+)_ group (figure 6A, B). Indeed MSC-CM contains other growth factors and soluble mediators that may also play a role in promoting lung repair and we cannot rule out the possibility that other factors may contribute the protective effects observed. MSC-CM_PPARβ/**δ**(+)_ did not significantly reduce the level of TNFα or IL-6 in the BALF of LPS-exposed mice (figure 6C-D). Administration of MSC-CM_PPARβ/**δ**(+)_ did not decrease weight loss or clinical score in LPS-exposed mice (figure 6E-F).

**Figure 6:**
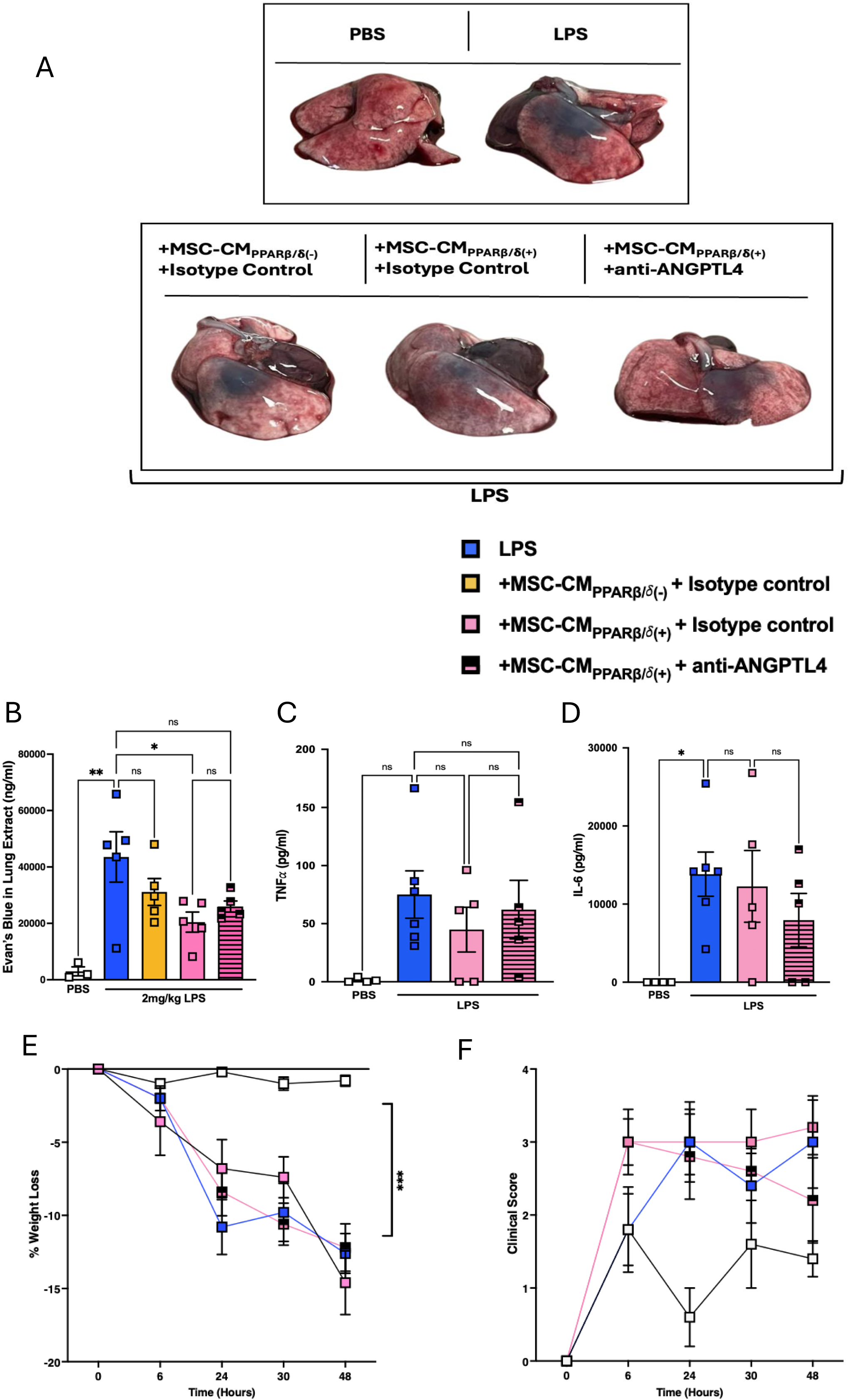
MSC-CM_PPARβ/**δ**(+)_ supports enhanced endothelial barrier function in ALI. Mice were exposed to 2mg/kg LPS IT, or PBS control. Some groups received MSC-CM_PPARβ/**δ**(-)_ or MSC-CM_PPARβ/**δ**(+)_ + isotype control or MSC-CM_PPARβ/**δ**(+)_ + anti-ANGPTL4. (A) Lungs were harvested and imaged after Evan’s Blue Dye administration highlighting increased vascular permeability (visible blue dye). (B) Quantification of the amount of Evan’s Blue dye leakage from the lungs of mice exposed to LPS +/- treatments of interest (n=5/group, or n=3 for PBS control group). Levels of pro-inflammatory cytokines in the bronchoalveolar lavage fluid (C) TNF⍺ and (D) IL-6 were assessed by ELISA (one-way ANOVA followed by Tukey’s post-hoc test, n=4-6/group). (E) % weigh loss and (F) Clinical scores (n=4-6/group) were recorded over the course of the model. Data points are a representation of individual mice. Data is presented as mean ± SEM; *p<0.05, **p<0.01.

### ARDS-licensed MSC-CM_PPARβ/**δ**(+)_ provide protection in ALI in an ANGPTL4 dependent manner

Given our findings that MSC-CM_PPARβ/**δ**(+)_ could enhance vascular and epithelial barrier function but could not decrease BALF pro-inflammatory cytokines or reduce weight loss and clinical score, we sought to investigate if we could increase the efficacy of MSC-CM. Interestingly exposure to ARDS patient serum, along with PPARβ/**δ** agonism, allowed for a 10-fold increased induction of ANGPTL4 in MSC-CM compared to PPARβ/**δ** agonist alone (figure 5K). Other proteins examined including IL-6, IL-8, VEGF, IL-1β and MIF (supplementary figure 4) were not increased in MSCs that had been exposed to ARDS patient serum, along with PPARβ/**δ** agonism. We hypothesised that ARDS-licensed MSC-CM_PPARβ/**δ**(+)_ would provide greater protection in ALI. In this case, the combination of ARDS serum and PPARβ/**δ** agonism led to enhanced therapeutic effects with reduction in TNFα and IL-6 in the BALF (figure 7A-B). Importantly, anti-ANGPTL4 reduced the anti-inflammatory effects mediated by the ARDS-licensed MSC-CM_PPARβ/**δ**(+)_ (figure 7A-B).

**Figure 7:**
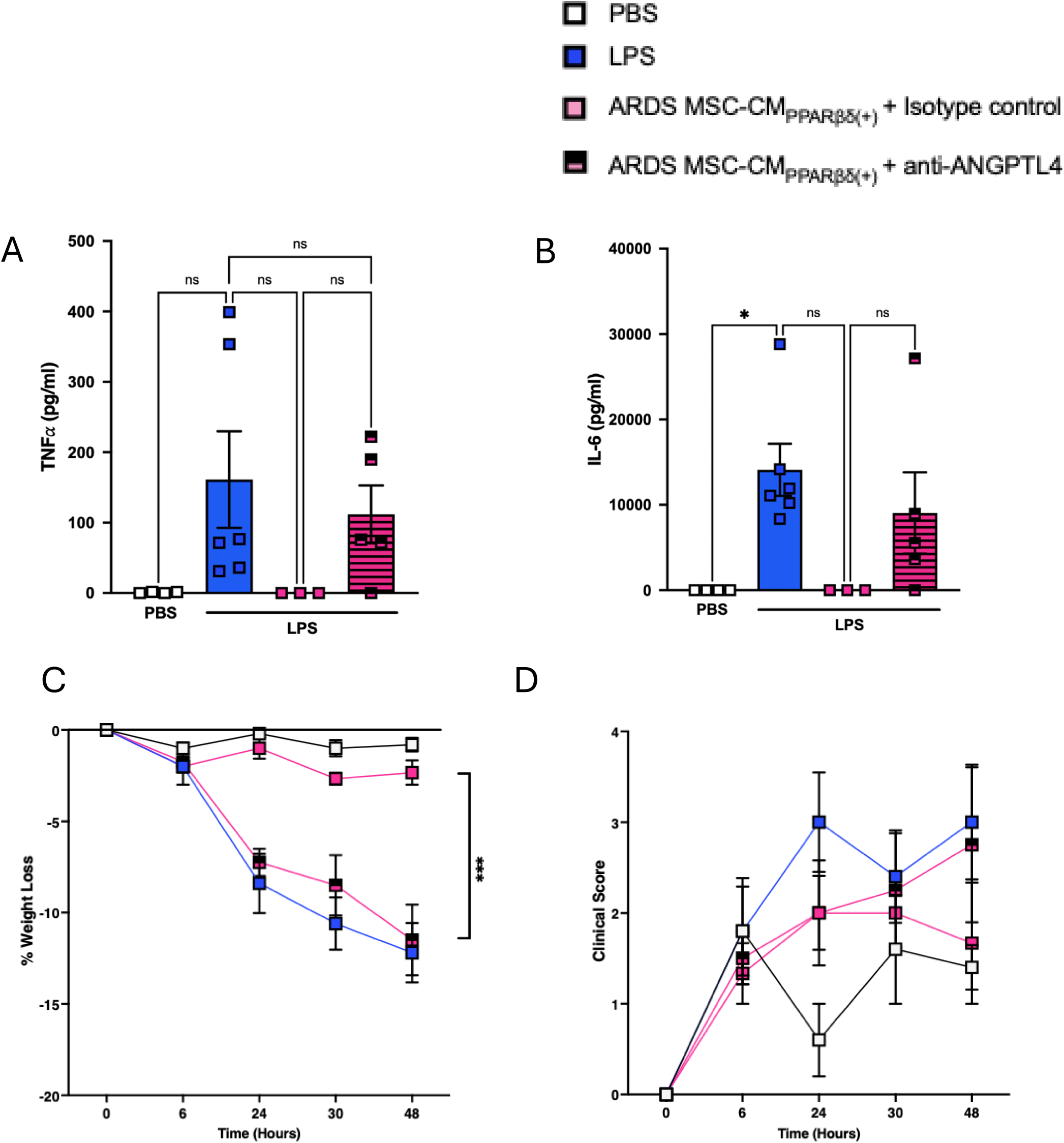
ARDS-licensed MSC-CM_PPARβ/**δ**(+)_ provide protection in ALI via ANGPTL4. Mice were exposed to 2mg/kg LPS IT, or PBS control. Some groups received ARDS licensed MSC-CM_PPARβ/**δ**(+)_ + isotype control or ARDS-licensed MSC-CM_PPARβ/**δ**(+)_ + anti-ANGPTL4. Levels of bronchoalveolar lavage fluid pro-inflammatory cytokines (A) TNF⍺ and (B) IL-6 were assessed by ELISA (one-way ANOVA followed by Tukey’s post-hoc test, n=3-6/group). (C) % weigh loss and (D) Clinical scores (n=3-6/group) were recorded over the course of the model. Data points represent individual mice. Data is presented as mean ± SEM; *p<0.05, ***p<0.001.

In line with this, there was a significant reduction in percentage weight-loss in mice exposed to ARDS-licensed MSC-CM_PPARβ/**δ**(+)_ (figure 7C), and again, this effect was abrogated in the presence of the anti-ANGPTL4-neutralising antibody (figure 7C). Interestingly, we saw a non-significant trend toward a reduced clinical score in mice exposed to ARDS-licensed MSC-CM_PPARβ/**δ**(+)_ (figure 7D). ANGPTL4-neutralisation appeared to abrogate this effect (figure 7D).

## DISCUSSION

There are two primary functions of MSCs in the context of cell therapy: immunomodulation and promotion of repair [1]. It is known that the environment can largely dictate MSC response [7, 11, 34]. In the context of ARDS, the complex patient microenvironment; containing inflammatory-agents, microorganisms and FFAs, can notably impact MSC functionality [10, 35]. We previously highlighted that higher levels of inflammation in the ARDS patient microenvironment can promote VEGF-driven repair [12]. Abreu *et al.* (2020) investigated the detrimental impact of fungal toxins present in the lung on MSC viability and showed a drastic increase in MSC cell-death in response to this microenvironmental element [7]. However, there has been little focus on how the FFAs in the ARDS patient microenvironment may impact MSCs [36]. FFAs are known natural ligands of PPARβ/**δ** [37–39]. Importantly, studies have already shown that modulation of this receptor can alter MSC functionality [17, 20, 21, 23, 25]. Given this, we wanted to investigate the functional impact of PPARβ/**δ**-modulation on both primary functions of MSCs; immunomodulation and/or pro-reparative capacity.

In the case of pro-reparative capacity, PPARβ/**δ**-activation enhanced MSC-CM mediated wound repair in CALU-3 lung epithelial cells (figure 1B, C). A time-lapse experiment, using a mitomycin C proliferation inhibitor, highlighted that this was likely a combination effect of both migration and proliferation (figure 1D, E). In the context of promoting tissue repair, MSC secretion of VEGF [40–43], and HGF [44, 45] among other factors have been shown to play key roles. However, PPARβ/**δ**-activation did not lead to differential expression of VEGF or HGF. Interestingly, ANGPTL4 was significantly increased in PPARβ/**δ** agonised MSCs. It has already been shown in several studies that ANGPTL4 can promote both migration and proliferation in endothelial cells [28, 29]. Ziveri *et al.* (2024) recently published a comprehensive study comparing the role of ANGPTL4 cytoprotection in skin and brain endothelial cells. ANGPTL4 in the brain endothelium protects against bacterial sepsis, and addition of ANGPTL4 to the skin endothelium, which naturally has lower levels of ANGPTL4, leads to protection against bacterial sepsis [27]. ANGPTL4 has also been shown to improve cardiac function in a mouse model of myocardial infarction [46]. Moreover, human MSCs produced increased levels of ANGPTL4 following co-culture with THP-1 macrophages *in vitro* [46]. Based on the identification of increased expression of ANGPTL4 in PPARβ/**δ** agonised MSCs in our RNA-seq data set, we used an anti-ANGPTL4 antibody to neutralise the ANGPTL4 in the MSC secretome, before addition to the CALU-3 lung epithelial cells (figure 4B-D). This abrogated the enhanced wound closure we had originally observed in our MSC-CM_PPARβ/**δ**(+)_ group.

We identified increased levels of PPARβ/**δ** ligands and co-activators in ARDS patient samples using publicly available datasets. Exposure to ARDS patient serum significantly increased ANGPTL4 levels produced by MSCs. A combination of MSC exposure to ARDS patient serum and a PPARβ/**δ** agonist further increased ANGPTL4 levels. This proves that MSCs are receptive to FFAs/other PPARβ/**δ** ligands presents in the ARDS microenvironment. This aligns to our previous study where we identified that human BM-MSCs produce significantly increased levels of ANGPTL4 in response to the FFA palmitate [14].

To enhance the clinical relevance of our findings, we showed that MSC-CM_PPARβ/**δ**(+)_ mediated protective effects in ALI significantly enhancing vascular and epithelial barrier integrity *in vivo*. Indeed, other studies have demonstrated that MSC-CM can mediate therapeutic effects *in vivo* in ALI models [47] including our own study [12]. Our previous study showed that MSC-CM generated from the equivalent of 5×10^4^ MSCs was ineffective in the treatment of ALI; but licensing 5×10^4^ MSCs with ARDS patient serum enhanced the therapeutic efficacy [12]. Using this approach of MSC-CM generated from the non-efficacious dose of 5×10^4^ MSCs, we investigated the potential for enhanced efficacy of MSC-CM generated from 5×10^4^ MSCs that had been exposed to a PPARβ/**δ** agonist or antagonist. MSC-CM_PPARβ/**δ**(+)_ provided enhanced efficacy over and above MSC-CM_PPARβ/**δ**(-)_ in a pre-clinical model of ALI. While there was a significant difference in the capacity for MSC-CM_PPARβ/**δ**(+)_ in enhancing barrier function, addition of the ANGPTL4-neutralising antibody failed to significantly impair this effect. This may be associated with growth factors or mediators other than ANGPTL4 present in the MSC-CM_PPARβ/**δ**(+)_ that may be contributing to the protective effects observed. Our study demonstrated that ARDS patient serum induced significantly higher levels of ANGPTL4 in MSC-CM and that combination with a PPARβ/**δ** agonist further increased it. While MSC-CM_PPARβ/**δ**(+)_ did not significantly reduce proinflammatory cytokines in the BALF, ARDS licensed MSC-CM_PPARβ/**δ**(+)_ reduced TNFα and IL-6 in the BALF of ALI mice. By combining the PPARβ/**δ** agonist and licensing with ARDS patient serum, we created MSC-CM with greater immunosuppressive capacity when compared to MSC-CM_PPARβ/**δ**(+)_ alone. This is likely associated with the ∼10-fold increase in ANGPTL4 levels when using the combined licensing approach as addition of the ANGPTL4 neutralising antibody reversed the trend. The limitation to these findings is the small numbers of mice (n=3/group) included in the study for the ARDS licensed MSC-CM_PPARβ/**δ**(+)_ treatment group. This was associated with a limited volume of ARDS patient serum available for the study.

Here we provide novel findings to demonstrate that human BM-MSCs are responsive to FFAs (ligands that bind PPARβ/**δ**) present in the ARDS patient environment, leading to significantly increased levels of ANGPTL4 production. For the first time, we show that MSC-derived ANGPTL4 plays an important role in promoting epithelial repair and enhancing vascular barrier integrity. As most published studies have focused on mouse MSCs, we have also addressed the gap in the knowledge of how PPARβ/**δ** influences human BM-MSC cytoprotective and immunosuppressive functions. In this study, focusing on ALI, we did not detect any significant positive or negative effects associated with PPARβ/**δ** antagonism in hBM-MSCs. However, PPARβ/**δ** agonism enhanced MSC-CM promotion of wound healing *in vitro* and vascular and barrier integrity *in vivo*. Furthermore, the combined approach of PPARβ/**δ** agonism and licensing with ARDS patient serum produced an MSC-CM that could suppress pro-inflammatory cytokines in the BALF of ALI mice.

## Declarations

This manuscript has been uploaded to Bioxriv as a pre-print.

## Author Contributions

C.T. performed research, data analysis, and study design and wrote the manuscript. M.D., S.R., E.V., E.J., S.B. and C.M. provided expertise and or performed research and data analysis. B.McN and J.G.L. provided patient samples and data for the study. C.C.D.S and J.G.L. contributed to study design. K.E. designed and supervised the study and wrote the manuscript. All authors approved the final manuscript.

## Ethical Approval and HPRA Compliance

Ethical approval was granted by the Maynooth University research ethics committee (BRESC-2022-2453953) granted 17^th^ February 2022 under the title: Cell based therapeutics for acute respiratory distress syndrome for the animal studies and by Galway University Hospitals Research Ethics Committee and by the Maynooth University research ethics committee (BSRESC-2022-2482563) granted 14^th^ October 2022 under the title: Human serum Sepsis/ARDS MSC. HPRA approval was granted under the project authorisation(s) AE19124/P031/P037.

RoosterBio (company where the human BM-MSCs were purchased) has confirmed that there was initial ethical approval for collection of human cells, and that the donors had signed informed consent. RoosterBio state “RoosterBio sources commercially available in vitro research only human bone marrow aspirate from qualified donors. All human bone marrow aspirate collections are from healthy adult consented donors. Collection protocols and the donor-informed consent document are approved by an Institutional Review Board (IRB)”

## Use of AI

The authors declare that they have not used AI-generated work in this manuscript.

## Data availability statement

The RNA-sequencing dataset generated in our lab is available through GEO accession no.: GSE281162. The other publicly available data sets are available online as outlined in the Methods section. The remaining data supporting the findings of this study are available upon request from the corresponding author.

## Acknowledgements

The authors are grateful to the ARDS patients who donated the samples used in this study. We would like to thank Rebecca Kerrigan, Deirdre Daly and Gillian O’Meara for the outstanding care of the animals used in this study. Schematic figures contained in this manuscript have been created using biorender.com.

