## Supplementary Tables and Figures for "The PPARβ/delta-induced mesenchymal stromal cell secretome has cytoprotective effects via ANGPTL4 in a pre-clinical model of acute lung inflammation"

| Gene name | Brand | Species | Forward Sequence | Reverse Sequence |
| --- | --- | --- | --- | --- |
| ***hprt*** | Sigma™ | Human | 5’ATAAGCCAGACTTTGTTGG | 5’ATAGGACTCCAGATGTTTCC |
| ***top2a*** | Sigma™ | Human | 5'TCACAAGCAAGAAATCCAAG | 5'ATCTTCATCTGACTCTTCCAG |
| ***angptl4*** | Sigma™ | Human | 5'AGGCAGAGTGGACTATTTG | 5'CCTCCATCTGAGGTCATC |
| ***pdk4*** | Sigma™ | Human | 5'CTTGGGAAAAGAAGACCTTAC | 5'GTGCAGTGGAGTATGTATAAC |
| ***ptgs2*** | Sigma™ | Human | 5'AAGCAGGCTAATACTGATAGG | 5'TGTTGAAAAGTAGTTCTGGG |
| ***il-6*** | Sigma™ | Human | 5'GCAGAAAAAGGCAAAGAATC | 5'CTACATTTGCCGAAGAGC |
| ***ido*** | Sigma™ | Human | 5'TTGTTCTCATTTCGTGATGG | 5'TACTTTGATTGCAGAAGCAG |
| ***ptges*** | Sigma™ | Human | 5'CAAAAACATCACTCCCTCTC | 5'AAAAGTCTGCATTCTTAGCC |
| ***tgfβ*** | Sigma™ | Human | 5'CCCACAACGAAATCTATGAC | 5'TGTATTTCTGGTACAGCTCC |

Supplementary table 1 (PRIMER SEQUENCES)

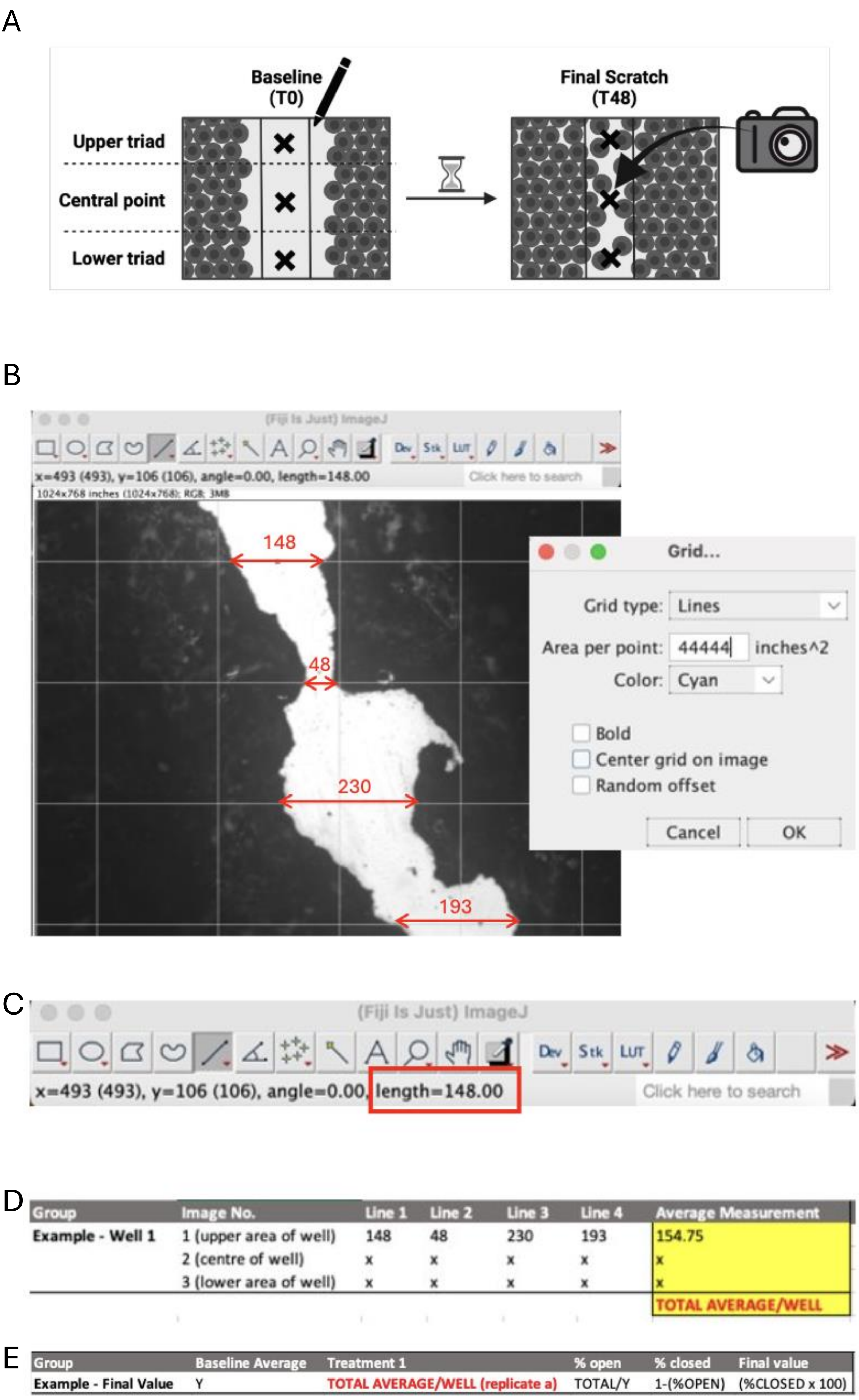


**Supplementary figure 1: Scratch assay analysis.** CALU-3s were seeded at a density of 5x10$^5$ and allowed to grow until confluent. A single scratch was made per well, and images were taken from a region in the upper triad, central point and lower triad before analysing with ImageJ/Fiji software.

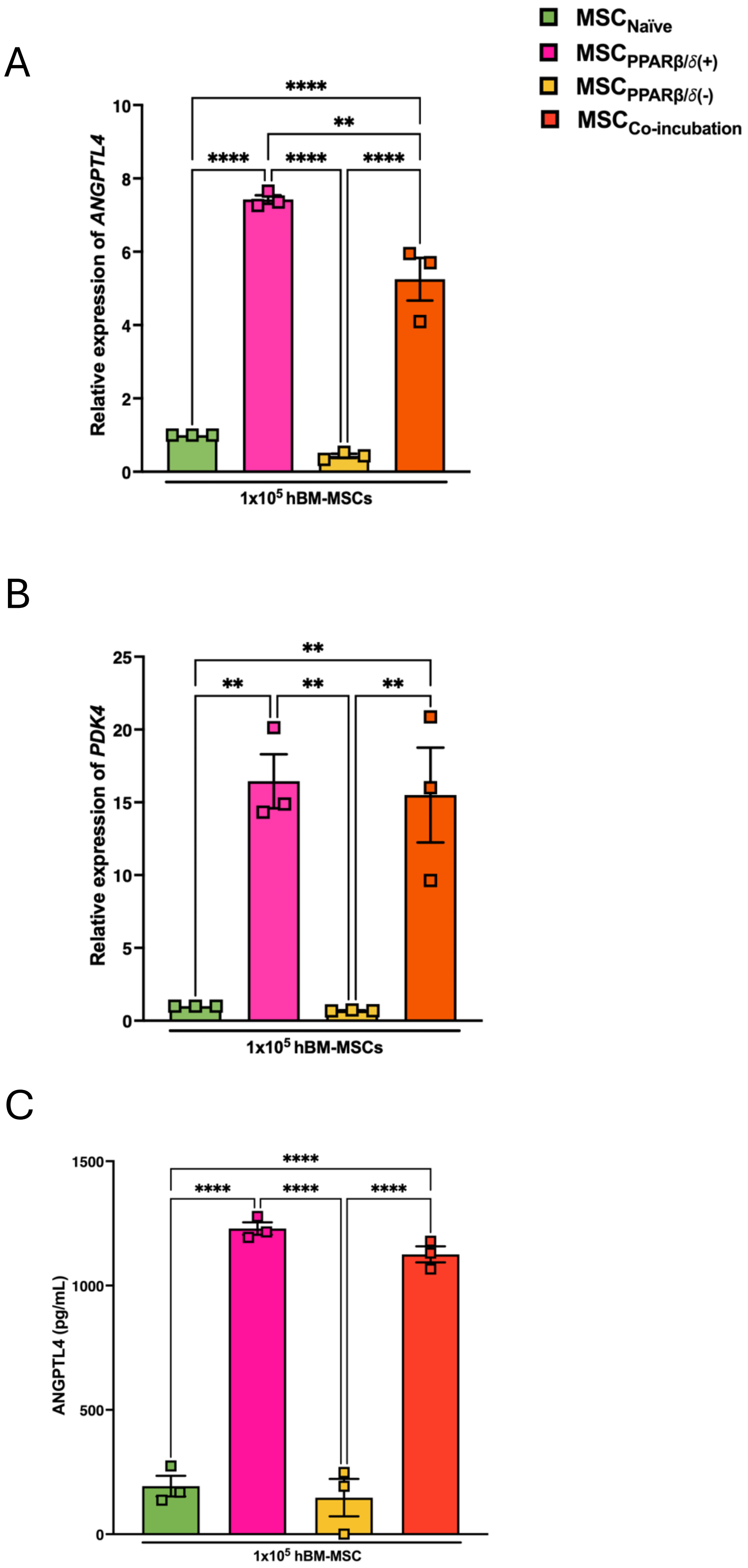


**Supplementary figure 2: Co-incubation of PPARβ/δ-agonised and antagonised hBM-MSCs.** hBM-MSCs were exposed to 1uM of a PPARβ/**δ** agonist *and* antagonist for 24hrs. Gene expression was assessed for (A) *angptl4* and (B) *pdk4* and protein expression was assessed for (C) ANGPTL4 (one-way ANOVA followed by Tukey's post-hoc test, n=3+). Replicates are a representation of individual hBM-MSC donors, with A and B previously presented in figure 2. Data is presented as mean ± SEM; **p<0.01, ***p<0.001, ****p<0.0001.

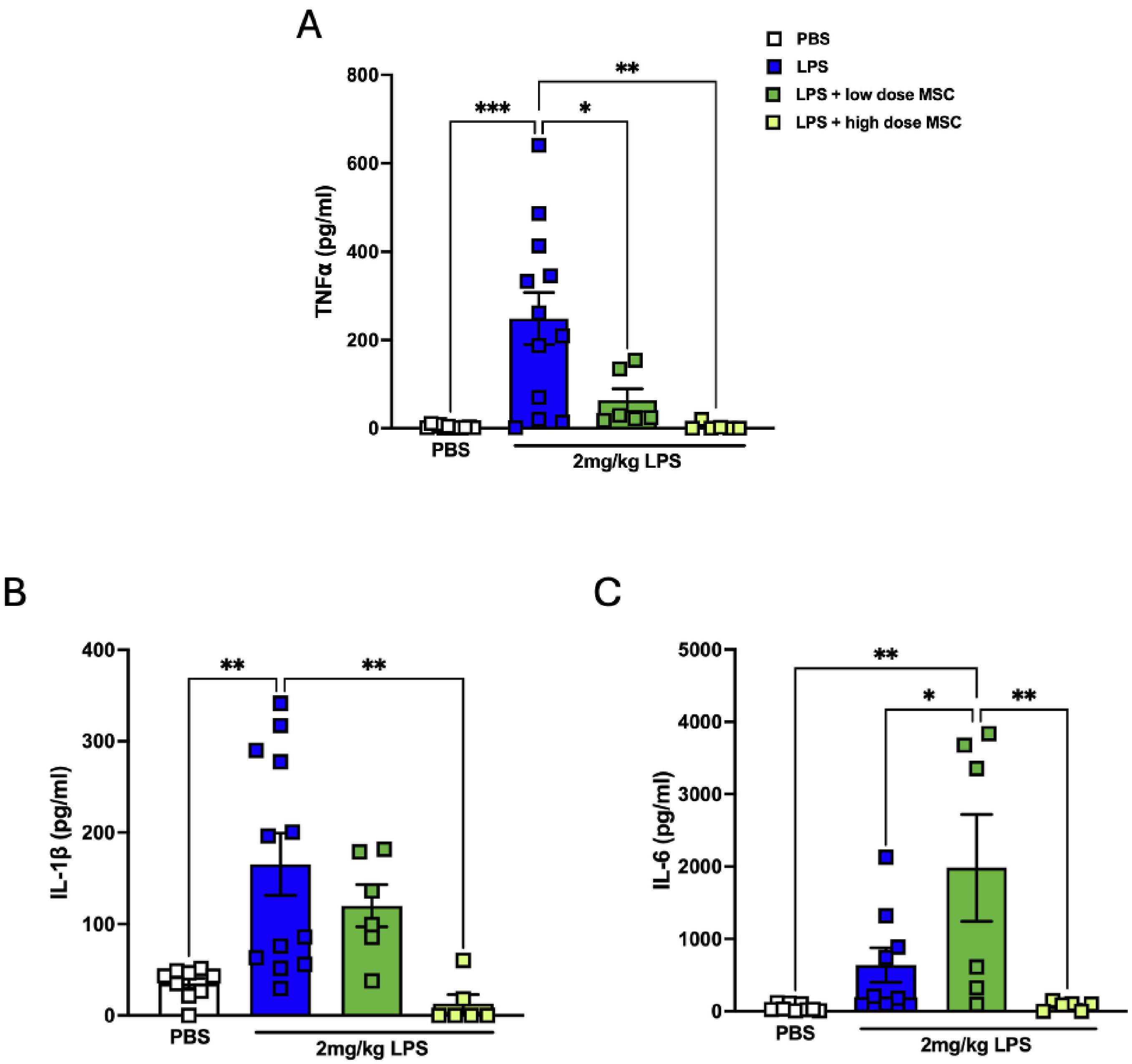


**Supplementary figure 3: ALI mouse model pilot using intratracheal administration and dose curve of hBM-MSCs.** Mice were exposed to 2mg/kg of LPS, or a PBS control, intratracheally and observed over 48hrs. They were further subjected to the administration of a low dose ($5x10^4$) or high dose ($1x10^5$) hBM-MSCs, intravenously. BALF pro-inflammatory cytokines (A) TNFα, (B) IL-1β and (C) IL-6 were measured by ELISA (one-way ANOVA followed by Tukey's post-hoc test, n=5-12). Replicates are a representation of individual mice. Data is presented as mean ± SEM; $*p<0.05$, $**p<0.01$, $***p<0.001$.

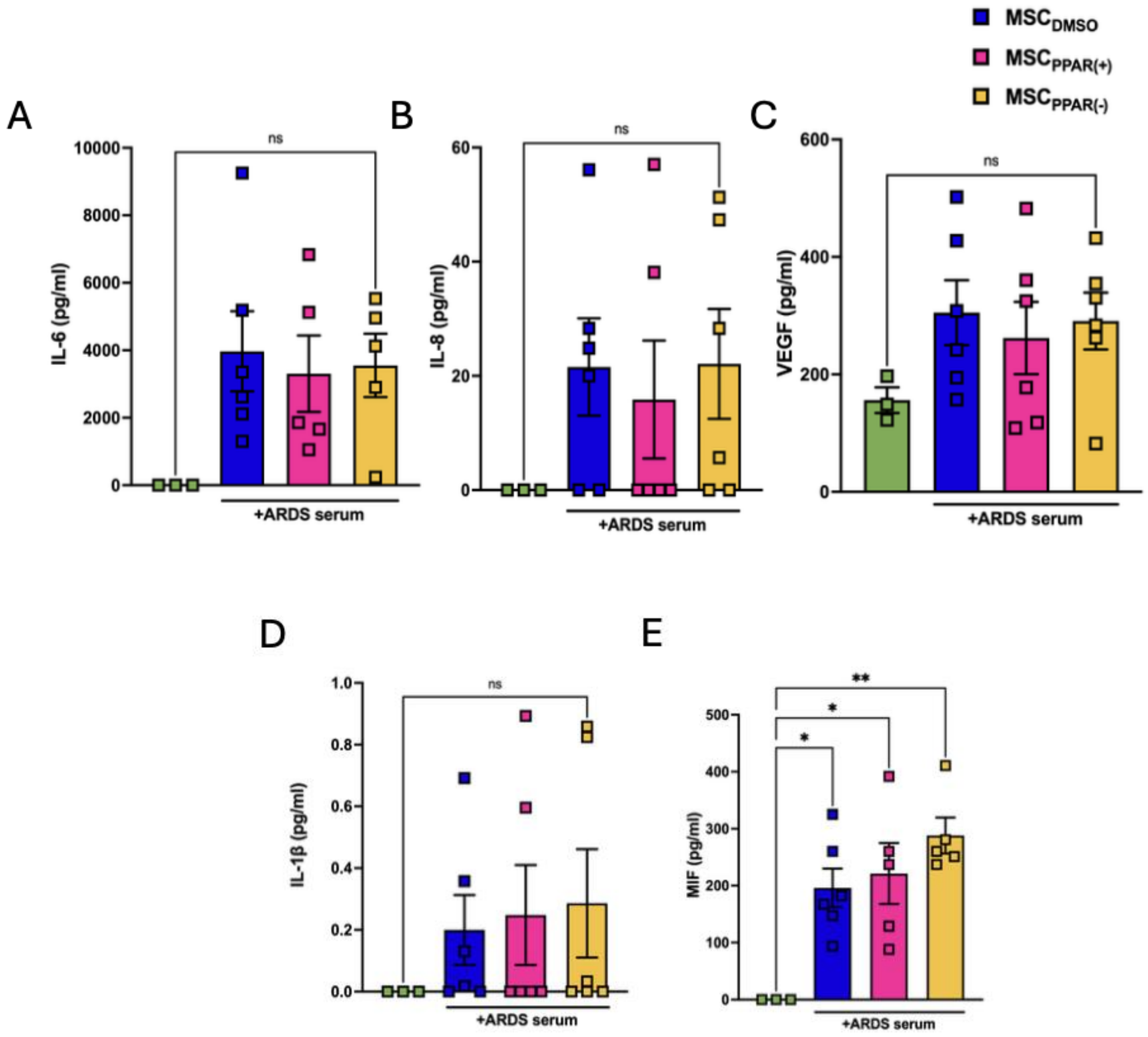


**Supplementary figure 4: PPARβ/δ(+/-) hBM-MSCs in response to ARDS patient serum.** hBM-MSCs were exposed to 1uM of a PPARβ/**δ** agonist or antagonist, and further exposed to ARDS patient serum. Protein expression was assessed for (A) IL-6, (B) IL-8, (C) VEGF, (D) IL-1β, (E) MIF (one-way ANOVA followed by Tukey's post-hoc test, n=3+). Replicates are a representation of individual hBM-MSC donors. Data is presented as mean ± SEM; *p<0.05, **p<0.01, ***p<0.001, ****p<0.0001.
